# Ensemble coding of crowd emotion: Differential hemispheric and visual stream contributions

**DOI:** 10.1101/101527

**Authors:** Hee Yeon Im, Daniel N. Albohn, Troy G. Steiner, Cody A. Cushing, Reginald B. Adams, Kestutis Kveraga

**Author notes:** Correspondence to: Hee Yeon Im, A.A. Martinos Center for Biomedical Imaging, Department of Radiology, Harvard Medical School, Massachusetts General Hospital, Charlestown, MA 02129,.

## Abstract

The visual system takes advantage of redundancies in the scene by extracting summary statistics from a set of items. Similarly, in many social situations where scrutinizing each individual’s expression is inefficient, human observers make snap judgments of crowds of people by reading “crowd emotion” to avoid danger (e.g., mass panic or violent mobs) or to seek help. However, how the brain accomplishes this feat remains unaddressed. Here we report a set of behavioral and fMRI studies in which participants made avoidance or approach decisions by choosing between two facial crowds presented in the left and right visual fields (LVF/RVF). Participants were most accurate for crowds containing task-relevant cues: avoiding angry crowds and approaching happy crowds. This effect was amplified by sex-linked facial cues (angry male/happy female crowds) and highly lateralized, with better recognition of the task-congruent facial crowd when presented in LVF. fMRI results showed that the dorsal visual stream was preferentially activated in crowd emotion processing, with intraparietal sulcus and superior frontal gyrus predicting behavioral crowd emotion efficiency, whereas the ventral visual stream showed greater involvement in individual face emotion processing, with fusiform cortex activity predicting the accuracy of decisions about individual face emotion. Our results shed new light on the distinction between global vs. local processing of face stimuli, revealing differential involvement of the left and right hemispheres and the dorsal and ventral pathways in reading crowd vs. individual emotion.

## Introduction

We routinely encounter groups of people at work, school, or social gatherings. In real-life situations, we often need to make quick decisions about which group of people to approach or avoid, and facial expressions of the group members play an important role in our judgments of their intent and predisposition. Because such decisions usually need to be made rapidly in many social situations, serial scrutiny of each individual’s facial expression is slow and becomes increasingly inefficient as the size of the crowd grows. Instead, extracting summary statistics (e.g., the average) through a process known as ensemble coding (for reviews, see Alvarez, 2011; Fischer & Whitney, 2011; Haberman, Harp, & Whitney, 2009; Haberman & Whitney, 2011) is a more efficient way to process an array of similar objects.

A large body of evidence has shown that the visual system can rapidly extract the average of multiple stimulus features such as orientation (Dakin & Watt, 1997; Parkes et al., 2001), size (Ariely, 2001; Chong & Treisman, 2003), and motion direction (Watamaniuk & Sekuler, 1992) of groups of objects in an array. Ensemble coding provides precise global representation (Alvarez, 2011; Ariely, 2001; Chong & Treisman, 2003; Halberda, Sires, & Feigenson, 2006), with little or no conscious perception (Alvarez & Oliva, 2008; Ariely, 2001; Choo & Franconeri, 2010; Corbett & Oriet, 2011; Parkes et al., 2001) or sampling of individual members in a set (Haberman & Whitney, 2010; Im & Halberda, 2013). Recent work has further shown that ensemble coding occurs for even more complex objects, such as averaging emotion from sets of faces (Fischer & Whitney, 2011; Haberman et al., 2009; Haberman & Whitney, 2007; Hubert-Wallander & Boynton, 2015; Ji, Chen, & Fu, 2014; Yang et al., 2013), facial identity (de Fockert & Wolfenstein, 2009; Haberman & Whitney, 2007; Leib et al., 2012; Leib et al., 2014; Neumann, Schweinberger, & Burton, 2013), as well as a crowd’s movements (Brunyé, Howe, & Mahoney, 2014; Sweeny, Haroz, & Whitney, 2013) and gaze direction (Florey et al., 2016; Sweeny & Whitney, 2014).

Face perception has great social importance, because emotional expressions forecast behavioral intentions of expressors (Adams et al., 2006; Horstmann, 2003; Marsh, Ambady, & Kleck, 2005) and govern observers’ fundamental social motivations accordingly (e.g., to approach or avoid; Elliot, 1999). To date, however, empirical work undertaken on ensemble perception of faces has largely concentrated on the efficiency and the fidelity of crowd perception (e.g., de Fockert & Wolfenstein, 2009; Fischer & Whitney, 2011; Haberman et al., 2009; Haberman & Whitney, 2007; Hubert-Wallander & Boynton, 2015; Ji et al., 2014; Leib et al., 2012; Leib et al., 2014; Neumann et al., 2013; Yang et al., 2013), but not on how this process is socially relevant. To our knowledge, no studies have examined how humans make speeded social decisions about which crowd of faces to approach or avoid, based on extracted ensemble features of facial crowds (e.g., crowd emotion). We often engage in such affective appraisals to enhance our social life (e.g., looking for a more approachable group of people to have a chat with at a cocktail party), and, occasionally, to avoid danger (e.g., rapidly inferring intent to commit violence from the facial expressions of a mob on the street to escape in time and seek help from another group that looks kinder).

Therefore, in the current study we aimed to characterize the behavioral and neural mechanisms of social decision making based on rapidly extracted crowd emotion from groups of faces. Behaviorally, our goals were to examine 1) how extracting crowd emotion from two groups of faces was modulated by observers’ social motivations (approach or avoidance) and whether this processing was significantly lateralized in the brain, and 2) how characteristics of facial crowds such as sex-linked identity cue and group size interact with the social motivations present when perceiving crowd emotion. These factors have received surprisingly little attention in the literature of ensemble perception of facial crowds, although they have been found to play a major role in affective processing elsewhere (e.g., Adams et al., 2006; Adams, Hess, & Kleck, 2015; Bargh, Chen, & Burrows, 1996; Craig, 2005; Davidson & Irwin, 1999; De Renzi, 1986; Fabes & Martin, 1991; Horstmann, 2003; Kanwisher et al., 1997; Marsh et al., 2005; Wada & Yamamoto, 2001).

Neurally, our goal was to examine the brain networks and pathways mediating ensemble perception of crowd emotion and compare it to the neural processes underlying extraction of emotion from, and choosing between, two individual faces. We focused on the dorsal and ventral visual streams, predicting their preferential involvement in processing crowd and individual emotion, respectively. The magnocellular (M) and parvocellular (P) pathways project primarily, but not exclusively, to the dorsal and ventral streams, respectively (Merigan & Maunsell, 1993; Sawatari & Callaway, 1996). The dorsal M-dominant pathway is suggested to support vision for action, non-conscious vision and detection of global and low-frequency information, whereas the ventral P-dominant pathway is suggested to support vision for perception, conscious vision, and analysis of local, high spatial frequency information (Freud, Plaut, & Behrmann, 2016; Goodale & Milner, 1992; Kveraga, Boshyan, & Bar, 2007; Kveraga, Ghuman, & Bar, 2007; Livingstone & Hubel, 1988; Milner & Goodale, 1995; 2008; Schiller & Logothetis, 1990; Thomas et al., 2012; Vuilleumier et al., 2003; Winston, Vuilleumier, & Dolan, 2003). Given such distinctive properties and functions of the dorsal and ventral pathways, we hypothesized that decisions on rapidly extracted global information of crowd emotion may rely on the dorsal pathway-dominant processing, whereas decisions that involve comparison between two individual emotional faces may rely on the ventral pathway-dominant processing. We tested this hypothesis by using both the whole brain and ROI analyses in the superior frontal gyrus (SFG) and the intraparietal sulcus (IPS) which have been implicated as conveying information via the M-pathway (Courtney et al., 1996; Courtney et al., 1998; Sala & Courtney, 2007; Takahashi, Ohki, & Kim, 2013; Wilson, Scalaidhe, & Goldman-Rakic, 1993) and the fusiform gyrus (FG) which is known to support P-pathway information (Denys et al., 2004; Grill-Spector & Malach, 2004; Haxby et al., 1991; Kanwisher et al., 1997; Purves et al., 2004; Takahashi et al., 2013; Taylor, Wiggett, & Downing, 2007). Since both anatomical and functional connectivity between IPS and SFG for M-pathway and the responsivity of FG for P-pathway have been shown to be predominant in the right hemisphere (Courtney et al., 1996; Takahashi et al., 2013), our ROI analyses focused on the right IPS, SFG, and FG (MNI coordinates are shown in the Method section).

To accomplish our goals, we conducted a set of behavioral and fMRI experiments in which participants viewed visual stimuli containing two groups of faces with varying emotional expressions (single faces were also examined for a direct comparison in the fMRI study), presented in the left and right visual hemifields. Participants had to choose one of the two crowds (or individual faces) as rapidly as possible, to indicate which one they would avoid or approach. Unlike the estimation task where the absolute value is judged, the answers and the ease of the decision in such comparison task would vary depending on the task goal. For example, the decision to choose to *approach* a happy crowd vs. an emotionally neutral crowd should be quite clear and explicit. However, the same comparison (happy vs. neutral) becomes more ambiguous and implicit when observers have to decide which crowd they would rather *avoid.* This paradigm allows us to examine the role of observers’ social motivation in comparing crowd emotion and its interaction with crowd size, sex-linked identity cues, and visual hemifield of presentation.

## Results

### Experiment 1: behavioral study

Participants viewed two crowds of faces (Figure 1B), one in the left visual field (LVF) and one in the right visual field (RVF) for 1 second. They were instructed to fixate on the center fixation cross and to make a key press as quickly and accurately as possible to indicate which group of faces they would rather avoid (Experiment 1A) or approach (Experiment 1B). The individual faces contained in each crowd were chosen from a set of 51 faces (Figure 1A) morphed from two highly intense, prototypical facial expressions (angry and happy) of the same person. The set contained six different identities (3 male and 3 female faces), taken from the Ekman face set (Ekman & Friesen, 1976). One visual hemifield always contained a crowd with varied expressions that were nonetheless emotionally neutral on the average (i.e., the particular mix of happier and angrier expressions was at the midpoint between happy and angry). The other visual hemifield contained a crowd that had a mix of expressions that was either happier or angrier on the average than the neutral crowd. Individual faces had all different emotional intensities, and half of individual faces were more intense in the neutral crowd than any expression in the emotional crowd. This is critical because it ensures that participants could not simply rely on finding the most intense (happy or angry) expression and base their decision that face. Instead, they had to choose the crowd to approach or avoid based on the *average* emotion from a crowd to perform the task correctly.

**Figure 1:**
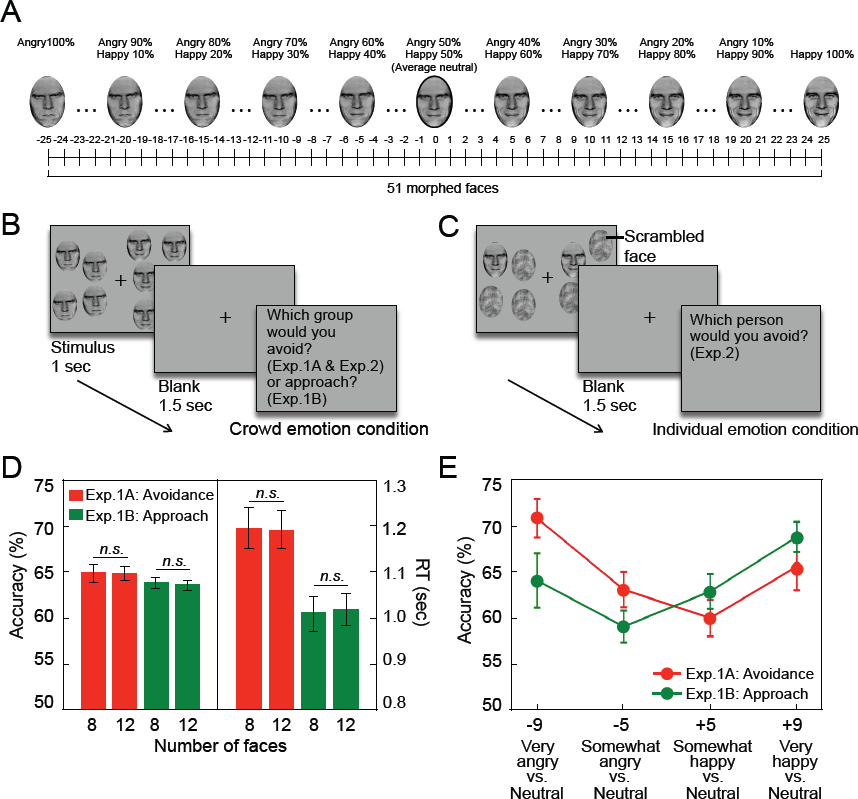
Sample face images, sample trials of crowd emotion and individual emotion conditions, and the results from Experiment 1. (A) Some examples of 51 morphed faces from two extremely happy and angry faces of the same person, with Face +25 in Emotional Unit (EU) being extremely happy, Face 0 being neutral, and Face -25 being extremely angry. (B) A sample trial of crowd emotion condition. (C) A sample trial of individual emotion condition (included in the fMRI study). (D) The effect of the number of faces on the accuracy and RT in Experiment 1A (avoidance task, red bars) and in Experiment 1B (approach task, green bars). The error bars indicate the standard error of the mean (SEM). (E) The effect of the similarity in average emotion between facial crowds on crowd emotion processing: Participants’ accuracies on Experiment 1A (avoidance task, red line) and Experiment 1B (approach task, green line) are plotted as a function of the emotional distance in EU between two facial crowds to be compared.

While participants reported the task as being rather difficult, they could reliably perform the task at levels well above chance. The overall accuracies for both Experiments 1A and 1B were significantly higher than chance (avoidance task: 64.88% vs. 50%; approach task: 63.72% vs. 50%, all *p’s* < 0.001), demonstrating that the participants were able to extract the average crowd emotion from the two groups of faces and choose appropriately which group they would rather avoid or approach. Although accuracies for the avoidance task vs. approach task were not significantly different (64.88% vs. 63.72%: *t*(40) = 1.330, *p* = 0.191), the mean response time (RT)^1^ was significantly slower for the avoidance task than for the approach task (1.17 vs. 0.98 second: *t*(40) = 2.156, *p* < 0.04, Cohen’s *d* = 0.666). Neither accuracy nor RT was affected by the size of the facial crowd (8 vs. 12 faces total) in the avoidance or approach task (Figure 1D), suggesting that extraction of crowd emotion does not require serial processing of each individual crowd member, but is processed in parallel. Because there was no effect of crowd size, we collapsed the data from the different crowd size conditions for further analyses.

#### Facilitation of task-congruent cues: avoiding angry and approaching happy crowds

In our morphing methods (Figure 1A), the emotional distance between the morphed faces could be quantified based on the arbitrary values of the emotional unit (EU) number, with zero being emotionally ambiguous (e.g., 50% happy and 50% angry), +25 being extremely happy (100% happy), and -25 being extremely angry (100% angry). Because the neutral crowd (EU of zero on average) was always presented on one side, the positive value of the emotional distance between the two crowds indicates that the other side to be compared contained a happier crowd than the neutral crowd (e.g., +9 vs. 0: very happy vs. neutral and +5 vs. 0: somewhat happy vs. neutral) and the negative value of the emotional distance indicates that the other side contained an angrier crowd than the neutral crowd (e.g., -9 vs. 0: very angry vs. neutral and -5 vs. 0: somewhat angry vs. neutral). Such separation proved to be effective in systematically manipulating the difficulty of the task (Figure 1E): In both the avoidance and approach tasks, accuracy increased when the emotional distance between the two crowds being compared increased (e.g., accuracy for ±9 EU was higher than for ±5 EU, all p’s < 0.005). A repeated-measures analysis of variance confirmed the significant main effect of the emotional distance (four levels: -9, -5, +5, and +9) on performance accuracy (avoidance Importantly, post-hoc contrast analyses revealed that participants were most accurate for the crowd emotion that was congruent with the task goal - whether to approach or to avoid. That is, subjects were most accurate when comparing a very angry crowd versus a neutral crowd (emotional distance of +9) during the *avoidance* task (Figure 1E, red line: F(1,20) = 12.659, *p* < 0.01, η_p_^2^ = 0.388) and when comparing a very happy crowd versus a neutral crowd (emotional distance of -9) during the *approach* task, than in comparing any other combinations (Figure 1E, green line: *F*(1,20) = 18.318, *p* < 0.01, η_p_^2^ = 0.504). The RTs were not significantly different for these conditions, although there was a trend toward faster RTs for the most task-congruent comparisons: very angry vs. neutral for avoidance and very happy vs. neutral for approach, compared to any other comparisons (Supplementary Information, SI.1A). These results suggest that observers were most accurate and efficient when they had to compare a task-congruent crowd emotion with a neutral crowd, with facilitated processing of angrier crowds for the avoidance task, and of happier crowds for the approach task. Thus, it appears that motivational information systematically modulates observers’ evaluation on crowd emotion: Comparing angry vs. neutral crowds allows for a *clear* and *explicit* decision in the avoidance task whereas comparing the same pair of crowd emotions requires more *ambiguous and implicit* decision in the approach task.

#### Hemispheric asymmetry for crowd emotion processing: explicit vs. implicit decisions

When participants judged which crowd they would avoid (Experiment 1A), choosing an angry over a neutral crowd was an explicit social decision directly relevant to the nature of the task. On the other hand, choosing a neutral over a happy crowd introduces ambiguity into the decision, because the neutral crowd does not contain an explicit social cue (although it is less friendly than a happy crowd). We observed a hemispheric asymmetry in participants’ accuracy for these explicit vs. implicit decisions on avoidance behavior based on crowd emotion when a task-congruent social cue (e.g., angry crowd) or an implicit cue (e.g., neutral crowd) was presented in LVF or RVF (illustrated in Figure 2A). The accuracy was higher when an explicit cue for avoidance task (e.g., an angry crowd) was presented in LVF than in RVF (red bars in Figure 2B). However, the pattern was completely reversed for an implicit cue (e.g., a neutral crowd). The accuracy was higher when an implicit cue was presented in RVF rather than in LVF (gray bars in Figure 2B). This observation was confirmed by a two-way repeated-measures ANOVA with a significant interaction between the visual field containing an emotional crowd (LVF vs. RVF) and the type of the cue contained in the emotional crowd (explicit vs. implicit: *F*(1,20) = 6.133, *p* < 0.05, η_p_^2^ = 0.235), although the main effect of the visual field (*F*(1,20) = 0.818, *p* = 0.376, η_p_^2^ = 0.039) and the main effect of the cue type (*F*(1,20) = 0.033, *p* = 0.858, = 0.002) were not significant. This result indicates hemispheric specialization in which LVF/RH presentations are superior for processing an explicit social cue (an angry crowd over neutral) whereas RVF/LH presentations are superior for processing a more implicit and ambiguous social cue (a neutral crowd over a happy one) during the avoidance task.

**Figure 2:**
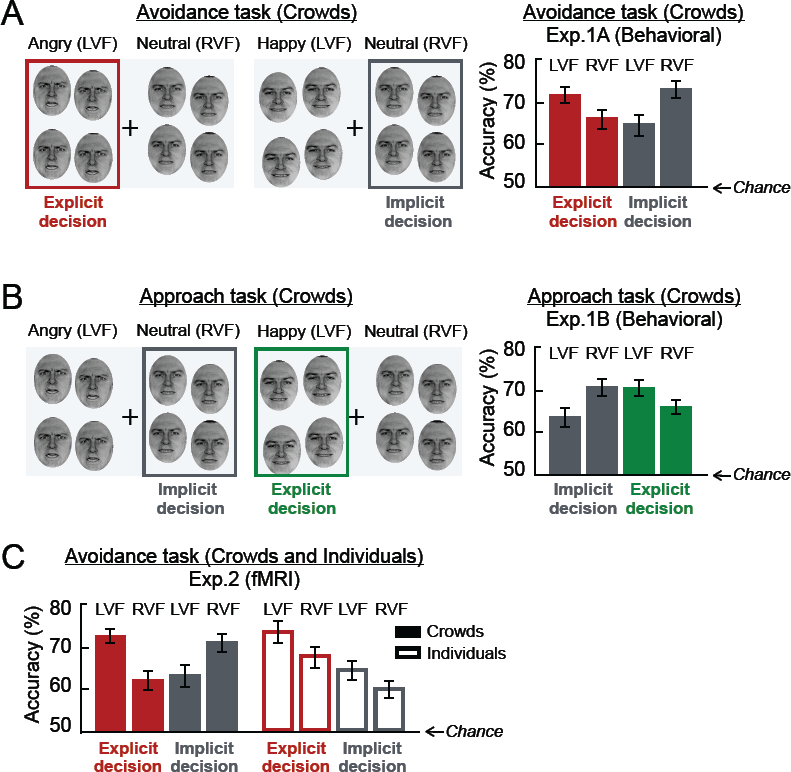
The task-goal dependent hemispheric asymmetry in the crowd emotion processing. (A) Participants’ accuracy for explicit and implicit decisions in the avoidance task (Experiment 1A), separately plotted for when the crowd to be chosen is presented in the LVF vs. RVF. When participants perform the avoidance task, choosing to avoid an angry crowd over neutral crowd becomes task-congruent and explicit decision (shown in red), whereas choosing a neutral crowd over a happy crowd is ambiguous and implicit decision (shown in gray). Participants’ accuracy was greater for explicit decision when the crowd to be chosen (e.g., angry) was presented in the LVF, but greater for implicit decision when the crowd to be chosen (e.g., neutral) was presented in the RVF. (B) Participants’ accuracy for explicit and implicit decisions in the approach task (Experiment 1B) in the LVF vs. RVF. Note that the valence of the task congruent crowd emotion is switched from angry to happy in the approach task. Participants’ accuracy was greater for explicit decision when the crowd to be chosen (e.g., happy, shown in green) was presented in the LVF, but greater for implicit decision when the crowd to be chosen (e.g., neutral, shown in gray) was presented in the RVF. (C) Participants accuracy for Experiment 2 (fMRI study). Accuracies both for crowd emotion and individual emotion conditions are plotted for the LVF and RVF, separately. As in Experiment 1, participants’ accuracy for crowd emotion condition was greater for the explicit decision (e.g., choosing an angry crowd) in the LVF and for the implicit decision in the RVF. However, patterns were different for individual condition: Accuracies were greater in the LVF both for the crowd emotion and individual emotion conditions, with the accuracy for the explicit decision (choosing an angry face over a neutral) greater than the implicit decision overall. The error bars indicate SEM.

For the approach task (Experiment 1B), on the other hand, participants had to choose which of the two crowds (e.g., Angry vs. Neutral or Happy vs. Neutral) they would rather approach. It is important to note that the emotional valence of the explicit and congruent social cue for the approach task is opposite to that for the avoidance task. For the approach task, choosing a happy over a neutral crowd is an explicit social decision whereas choosing a neutral over an angry crowd is a more ambiguous decision (illustrated in Figure 2D). Despite the emotional valence of a task-congruent social cue being flipped (e.g., angry for avoidance and happy for approach task), we again found a consistent pattern of hemispheric asymmetry for explicit and implicit social decision for approach decisions: the LVF/RH was superior for the *explicit* social cue (i.e., happy vs. neutral), while the RVF/LH was superior for the *implicit* social cue (i.e., neutral vs. angry). The accuracy was higher when an explicit, task-congruent cue (e.g., a happy crowd) was presented in LVF/RH than in RVF/LH (green bars in Figure 2E). As in the avoidance task (Experiment 1A), the accuracy was higher when an implicit cue (e.g., a neutral crowd) was presented in RVF/LH than in LVF/RH (gray bars in Figure 2E). This interaction between the visual field containing the crowd to be chosen and the type of social cue (explicit vs. implicit) conveyed by the crowd was confirmed by a two-way repeated measures ANOVA (F(1,20) = 5.447, *p* < 0.05, η_p_^2^ = 0.232).

Two-way ANOVAs (factors: visual field and type of cue) of participants’ mean RT for the avoidance and approach tasks (Figures shown in SI.1B) revealed that the main effect of type of cue (explicit vs. implicit) was significant (avoidance task: F(1,20) = 20.687, *p* < 0.001, η_p_^2^ = 0.506 and approach task: F(1,20) = 16.541, *p* < 0.001, η_p_^2^ = 0.479), although the main effect of visual field (avoidance task: F(1,20) = 0.470, *p* = 0.470, η_p_^2^ = 0.026 and approach task: F(1,20) = 1.317, *p* = 0.266, η_p_^2^ = 0.068) and the interaction (avoidance task: F(1,20) = 2.888, *p* = 0.105, η_p_^2^ = 0.126 and approach task: F(1,20) = 0.298, *p* = 0.592, η_p_^2^ = 0.016) were not significant. These RT results indicate faster processing of crowds containing clear cues than of crowds containing ambiguous cues, for both the avoidance and approach tasks. Furthermore, our RT results suggest that the differences in accuracy observed in all our studies are not due to a speed-accuracy trade-off.

Together, our results suggest that an LVF presentation, which is initially perceived by the right hemisphere, is dominant for processing an explicit, goal-congruent social cue in the context of extracting crowd emotion, preferring aversive and positive stimuli during avoidance and approach decisions, respectively, whereas an RVF/LH presentation dominant the processing of more ambiguous and implicit social cues. Unlike the traditional framework of face processing, which posits right hemispheric lateralization for aversive or negative face stimuli, and left hemispheric preference for positive, approach-evoking stimuli (Davidson, 1992, 1995; Davidson & Irwin, 1999; Silberman & Weingartner, 1986), our data suggest that the pattern of the hemispheric lateralization, at least for reading crowd emotion, is modulated in a more flexible manner depending on the task goal and decision uncertainty, rather than being based purely on stimulus valence.

#### Sex-specific identity cues that modulate crowd emotion perception

Because previous findings of individual face perception have documented that female-and male-specific facial features are perceptually confounded with happy and angry expressions, respectively (Adams et al., 2015; Becker et al., 2007), we examined whether processing of crowd emotion was also modulated by sex-specific facial identity cues. We compared the accuracy for male and female facial crowd stimuli (illustrated in Figure 3A) in the different tasks. Figures 3B and 3C show the accuracy on the avoidance task and approach task, plotted separately by the valence of emotional face images (happy and angry) to be compared to a neutral crowd and the sex of the face images (male vs. female). We found that participants’ responses were more accurate for happy than angry female crowds, while the opposite was true for angry male crowds, which were more accurately recognized than happy male crowds. The two-way repeated-measures ANOVA confirmed this observation. We observed a significant interaction between the sex and the emotion of the face images both in the avoidance task (F(1,20) = 4.908, *p* < 0.04, η_p_^2^ = 0.197) and in the approach task (F(1,20) = 4.678, *p* < 0.05, η_p_^2^ = 0.190). Although the main effects of the stimulus sex (F(1,20) = 2.984, *p* = 0.100) or of the emotional valence of the face images (F(1,20) = 0.112, *p* = 0.741) were not significant in the avoidance task (all *p’s* > 0.160), we also found the significant main effect of the stimulus sex in the approach task (better accuracy for female crowds: F(1,20) = 4.966, *p* < 0.04, η_p_^2^ = 0.199), but not the main effect of the emotional valence (F(1,20) = 0.588, *p* = 0.981). Together, these results suggest that integration of crowd emotion from emotional faces is also influenced by sex-specific identity cues.

**Figure 3:**
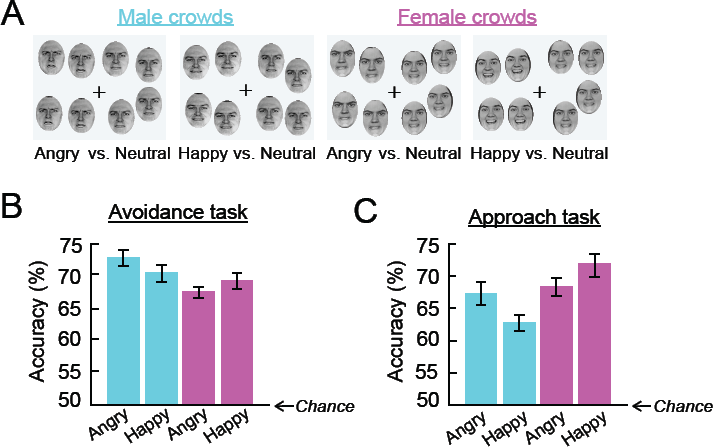
The effect of the sex-specific identity cue of facial crowds on crowd emotion perception. (A) Sample crowd stimuli for male (in cyan) and female (in magenta) crowds. (B) Participants’ accuracy for the avoidance task (Experiment 1A) for sex of facial crowds (male crowds vs. female crowds) and for the emotional valence of an emotional crowd (Angry vs. Happy). Angry female crowds were identified most accurately in the avoidance task. (B) Participants’ accuracy for the approach task (Experiment 1B). In the avoidance task, happy female crowds were identified most accurately.

Comparing the differing task demands for avoidance and for approach, we also observed a modulation by task demands. Contrast analyses revealed that participants were most accurate for comparing an angry male crowd vs. a neutral male crowd during the avoidance task), suggesting that facial anger and masculine features both conveyed threat cues and interacted to facilitate decisions to avoid a crowd. Conversely, participants were most accurate in comparing a happy female crowd vs. a neutral female crowd during the approach task. Although the sex of the faces in our crowd stimuli modulated the perception of crowd emotion, we found that the sex of the participants did not influence perception of crowd emotion either in the avoidance or the approach task (SI.2).

Finally, we examined whether increased variability in facial identities per se interferes with reading of crowd emotion and tested the robustness of our effects in two additional replication and extensions experiments in which participants were presented with crowd stimuli containing a mix of different identities (SI.3). We replicated the results of Experiments 1A and 1B using facial crowds with mixed identities and with new cohorts of participants (SI.4), confirming the robustness of our findings of task-goal dependent modulation and hemispheric asymmetry for explicit and implicit decisions on crowd emotion.

### Experiment 2: fMRI study

In the fMRI study, we scanned 30 participants, using only the avoidance task because of time and budgetary constraints. Participants were presented with stimuli containing either two facial crowds (Figure 1B) or two single faces presented in a crowd of scrambled masks (Figure 1C). Participants were asked to choose rapidly which of the two facial crowds (crowd emotion condition) or which of the two single faces (individual emotion condition) they would rather avoid, using an event-related design with crowd emotion and individual faces conditions randomly intermixed (See Methods for more details). We compared the patterns of brain activation when participants chose to avoid one of two crowds or one of two individual faces. If the processing of crowd emotion relies on the same mechanism that mediates single face perception, we would observe activations of the same brain network during the processing of crowd emotion, but perhaps to a larger degree and larger extent than during single face comparisons, given the greater complexity of the stimulus and difficulty of the crowd emotion task. Alternatively, if the processing of crowd emotion and of individual face emotion relies on qualitatively distinct processes, specifically mediated by dorsal and ventral visual pathways as we hypothesized, we would expect to observe differential brain activations in distinct sets of brain areas in dorsal and ventral visual pathways.

### Behavioral results

We first confirmed that the behavioral results on the crowd emotion condition from our fMRI study again replicated our behavioral results in Experiment 1A and in the replication and extension experiments. The participants’ overall accuracy for crowd emotion condition in the fMRI study was 63.16%, not significantly different from that we observed from Experiment 1A (64.88%; t(48) = -1.468, *p* = 0.149). Critically, we again replicated the laterality effect on the avoidance task we found in Experiment 1A (see Figures 2B) and in the replication and extension experiments (see SI.4). The participants’ accuracy was higher when faced with an explicit decision, task-congruent angry crowd was presented in LVF than in RVF, and when an implicit, neutral crowd was presented in RVF than in LVF (Figure 2C). This was confirmed by a significant interaction between the visual field of presentation (LVF vs. RVF) and the type of cue contained in an emotional crowd (explicit vs. implicit) in a two-way repeated measures ANOVA (F(1,28) = 6.357, *p* < 0.05; main effects were not significant). For the individual emotion condition, the overall accuracy was 65.92%, which was slightly but not significantly, higher than that for crowd emotion condition (*t*(58) = -1.491, *p* = 0.106). We also observed no difference in the RT for the crowd emotion condition vs. the individual emotion condition (*t*(58) = 0.318 *p* = 0.751 ). Even though only two faces were presented, and thus there was no need to extract the average crowd emotion, the level of accuracy and the response time for comparing two individual faces was similar to that for comparing two facial crowds. These results confirm that the difference in our fMRI findings comparing crowd emotion vs. individual emotion conditions is not due to a difference in task difficulty, but reflects qualitative differences in neural processing patterns and substrates.

Finally, we observed different patterns of hemispheric lateralization for the individual emotion condition and the crowd emotion condition (Figure 2G). Unlike the crowd emotion condition, in which we found better accuracy for explicit threat cues in the LVF and ambiguous threat cues in the RVF, individual emotion condition showed that both explicit and implicit threat cues were more accurately identified when presented to the LVF/RH than to the RVF/LH. In addition, an explicit threat cue (an angry face) was identified more accurately both in the LVF/RH and in RVF/LH presentations. A two-way repeated-measures ANOVA confirmed this observation: The main effects of the visual field of presentation (LVF vs. RVF: *F*(1,28) = 8.193, *p* < 0.01) and of the type of the threat cue (explicit vs. implicit: *F*(1,28) = 18.511, *p* < 0.001) were significant, but the interaction between the visual field of presentation and emotional valence was not significant (*F*(1,28) = 1.702, *p* = 0.203). This pattern of hemispheric lateralization found in the individual emotion condition is consistent with the previous findings suggesting that affective face processing in general is right-lateralized, with more marked laterality effects for the negative valence (Becker et al., 2007; Borod et al., 1998; Davidson & Irwin, 1999; Silberman & Weingartner, 1986). Together, our behavioral data from the fMRI study replicate our main findings from Experiment 1A and provide further evidence that perception of crowd emotion and individual emotion engage different patterns of hemispheric specialization.

### fMRI results

#### Distinct neural substrates for crowd emotion vs. individual emotion processing

Our goal in the fMRI experiment was first to characterize the neural substrates involved in participants’ avoidance decision between two facial crowds vs. those mediating decisions between two individual faces. Figure 4A shows the brain regions activated when participants were comparing two crowds (labeled in red) vs. comparing two individual faces (labeled in blue) and vice versa (The complete list of activations is reported in the Table 1). We observed that comparing two facial crowds with the task goal of avoidance showed greater cortical activations in the occipital (Brodmann area (BA) 19), parietal, and frontal regions along the dorsal stream (e.g., IPS, dmPFC, MFG, SFG, and OFC) in both hemispheres. On the other hand, deciding which of two individual faces to avoid evoked greater activation in the regions along the ventral stream (FG, LG, PHC, and RSC), PC/PCC and vmPFC.

**Figure 4:**
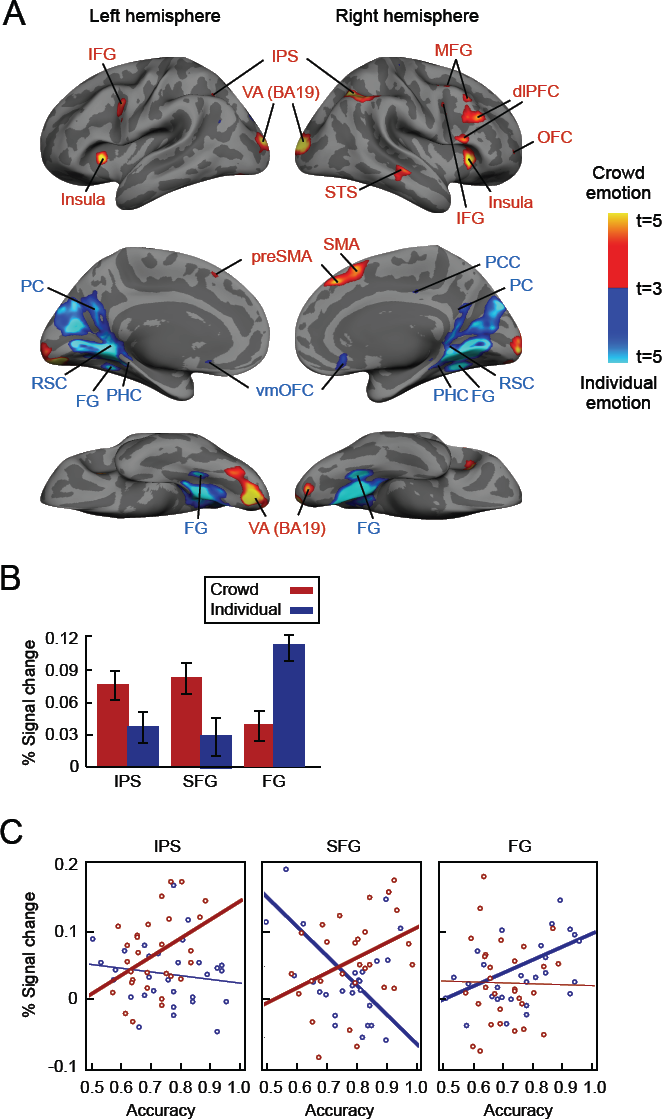
Distinct neural pathways preferentially involved in dorsal and ventral visual pathways for crowd emotion and individual emotion processing, respectively. (A) The brain areas that showed greater activation when participants were making avoidance decision by comparing two crowds are shown in red and the brain areas that showed greater activation for comparing two single faces are shown in blue. (B) The percent signal change of our ROI’s (IPS, SFG, and FG) when participants were making avoidance decision by comparing two crowds (red bars) and two single faces (blue bars). (C) The correlation between the percent signal change and the participants’ accuracy for crowd emotion condition (red dots) and for individual emotion condition (in blue dots), with overlaid linear regression lines. Thick regression lines indicate statistically significant correlation between the individual participants’ accuracy and the percent signal change of each ROI.

**Table 1:** The list of regions of increased activation associated showing greater activation for crowd emotion condition vs. individual condition.

| Activation location | MNI Coordinates |  |  |  |  |
| --- | --- | --- | --- | --- | --- |
| **Crowd emotion > Individual emotion** | **x** | **y** | **z** | **t-value** | **Extent** |
| L Visual cortex (BA18) | -15 | -88 | -14 | 7.872 | 1524 |
| R Visual cortex (BA18) | 18 | -100 | 12 | 6.174 | 692 |
| L Visual cortex (BA19) | -21 | -97 | 14 | 5.241 | 1524 |
|  | -33 | -76 | -16 | 4.730 | 1524 |
| L Anterior Insula | -33 | 20 | 4 | 5.973 | 146 |
| R Anterior Insula | 30 | 26 | 2 | 5.890 | 2401 |
| L Middle frontal gyrus | -42 | 29 | 36 | 3.072 | 26 |
| R Middle frontal gyrus | 36 | 32 | 46 | 5.826 | 2401 |
| L Superior frontal gyrus | -24 | 11 | 64 | 3.110 | 10 |
|  | -27 | -1 | 70 | 2.956 | 11 |
| R Superior frontal gyrus | 33 | 8 | 64 | 5.664 | 2401 |
| L Intraparietal sulcus | -39 | -49 | 38 | 4.731 | 353 |
| R Intraparietal Sulcus | 39 | -58 | 48 | 5.665 | 889 |
| R Supplementary Motor Area | 9 | 32 | 48 | 5.187 | 430 |
| L Superior parietal lobule | -30 | -67 | 48 | 3.875 | 353 |
| R Superior temporal sulcus | 48 | -25 | -10 | 4.661 | 242 |
|  | 39 | -34 | 4 | 3.357 | 10 |
| L Premotor cortex | -39 | 5 | 36 | 4.653 | 268 |
|  | -30 | 2 | 46 | 3.129 | 13 |
| L Thalamus | -6 | -10 | 20 | 3.797 | 293 |
| R Thalamus | 9 | 2 | 10 | 3.727 | 293 |
| L Inferior frontal gyrus | -27 | 26 | 22 | 3.499 | 12 |
| L Anterior prefrontal cortex | -42 | 59 | 8 | 3.236 | 48 |
| L Caudate | -12 | 5 | 10 | 3.003 | 5 |
| R Caudate | 12 | 23 | 8 | 3.148 | 9 |
| R Cerebellum | 33 | -52 | -44 | 3.282 | 8 |
| **Individual emotion > Crowd emotion** | **x** | **y** | **z** | **t-value** | **Extent** |
| R Visual cortex (BA19) | 12 | -64 | -8 | 8.054 | 2557 |
| L Visual cortex (BA19) | -18 | -55 | -6 | 6.948 | 469 |
| R Visual cortex (BA19) | 48 | -79 | 18 | 4.276 | 169 |
| L Primary visual cortex (BA17) | -3 | -76 | 10 | 6.188 | 5120 |
| L Fusiform gyrus | -24 | -49 | -12 | 6.191 | 4557 |
| R Fusiform gyrus | 21 | -40 | -14 | 5.283 | 4557 |
| L Parahippocampal cortex | -27 | -31 | -14 | 4.694 | 469 |
| R Parahippocampal cortex | 21 | -37 | -14 | 5.252 | 589 |
| L Temporal pole | 51 | 23 | -38 | 4.425 | 26 |
| R Posterior cingulate cortex | 12 | -25 | 42 | 4.364 | 64 |
| L Angular gyrus | -42 | -67 | 24 | 3.700 | 142 |
| L Ventromedial orbitofrontal cortex | -9 | 44 | 8 | 3.675 | 303 |
|  | -21 | 14 | -22 | 3.456 | 19 |
| R Ventromedial orbitofrontal cortex | 6 | 26 | -8 | 4.176 | 232 |
|  | 9 | 47 | -8 | 3.571 | 303 |
| L Precuneus | -12 | -28 | 40 | 3.627 | 55 |
| R Precuneus | 6 | -52 | 64 | 3.100 | 6 |
| L Posterior insula | -39 | -16 | 0 | 3.440 | 17 |
| R Hippocampus | 27 | -13 | -22 | 3.144 | 7 |

We further examined responses in our regions of interest (IPS, SFG, and FG). We chose these ROIs based on the prior work that showed involvement of SFG in dorsal stream processing and its intrastream functional connectivity with IPS (e.g., Courtney et al., 1996; Courtney et al., 1998; Sala & Coutney, 2007; Takahashi et al., 2013; Wilson et al., 1993) and the work showing the major involvement of FG in ventral stream information processing (e.g., Denys et al., 2004; Grill-Spector & Malach, 2004; Haxby et al., 1991; Kanwisher et al., 1997; Purves et al., 2004; Takahashi et al., 2013; Taylor et al., 2007). These regions were functionally restricted based on an *unbiased* contrast of all visual conditions minus baseline (average activation of voxels) using random effects models (height: *p* < 0.01, uncorrected; extent: 5 voxels), within the anatomical label for each ROI (obtained by the anatomical parcellation of the normalized brain; Tzourio-Mazoyer et al., 2002). As shown in Figure 4B, we observed that IPS and SFG showed greater activation for the crowd emotion condition (red bars in Figure 4B) than the individual emotion condition (blue bars in Figure 4B), whereas FG showed greater activation for the individual emotion condition than crowd emotion condition. This observation was confirmed by independent samples t-tests (two-tailed), conducted separately for each ROI (crowd emotion > individual emotion: IPS: t(58) = 2.203, p < 0.04 and SFG: t(58) = 2.134, p < 0.04; individual emotion > crowd emotion: t(58) = 2.027, p < 0.05). These results suggest that the regions in the dorsal visual pathway (e.g., IPS and SFG) and in the ventral visual pathway (e.g., FG) show differential responsivity to the crowd emotion and the individual emotion comparisons, respectively. Furthermore, we found that the activity in these ROIs differentially predicted the participants’ behavioral accuracy for the crowd emotion and individual emotion conditions. As shown in Figure 4C, the activity of IPS and SFG positively correlated with participants’ accuracy for the crowd emotion condition (IPS: *r* = 0.462, *p* < 0.01; SFG: *r* = 0.454, *p* < 0.02), whereas the activity of FG positively correlated with the accuracy for the individual emotion condition (FG: *r* = 0.498, *p* < 0.01). Together, our fMRI results provide first evidence for the differential contributions of the dorsal and ventral visual pathways to crowd emotion and individual emotion processing, respectively.

## Discussion

The goal of this study was to characterize the functional and neural mechanisms that support crowd emotion processing. We had four main findings: 1) Goal-dependent hemispheric asymmetry in which presenting a clear task-congruent cue (e.g., angry crowd for avoidance, and a happy crowd for approach) to the left visual field/right hemisphere led to higher accuracy, whereas an RVF/LH presentation was superior for implicit decisions involving an ambiguous cue (e.g., neutral crowd vs. one displaying a task-incongruent emotion); 2) Higher accuracy overall in identifying facial crowds containing task-congruent cues (e.g., angry crowd to avoid, happy crowd to approach); 3) Accuracy was further improved by sex-linked identity cues congruent with the task goal (e.g., angry male crowds to avoid and happy female crowds to approach); and 4) The dorsal visual stream was preferentially activated in crowd emotion processing, with IPS and SFG predicting behavioral crowd emotion efficiency, whereas the ventral visual stream showed greater involvement in individual face emotion processing, with fusiform cortex activity predicting the accuracy of decisions about individual face emotion.

The goal-dependent modulation of crowd emotion processing suggests that the mechanism underlying the reading of crowd emotion is highly flexible and adaptive, allowing perceivers to focus most keenly on desired outcomes in dynamic social contexts (e.g., to avoid unfriendly crowds or to approach friendly ones). It is worth noting that neither the stimulus display nor the response characteristics changed between avoidance and approach tasks: the only difference was the decision (approach or avoid) that was mapped to the response. The same visual stimuli containing facial crowds appear to be biased differently depending on whether the task goal was to avoid or to approach. Stimulus gender also interacted with the processing of crowd emotion, in a manner relevant to the current goal. Such visual integration of compound social cues (e.g., gender, emotion, race, eye gaze, body language, etc.) has been well incorporated into the theories of mechanisms underlying single face perception (for review, see: Adams & Kveraga, 2015). However the roles of these compound social cues in ensemble coding of facial crowds have not been examined. The current study provides new evidence that intrinsic (observers’ motivation) and extrinsic (e.g., emotional expressions and sex of the crowds) factors also mutually facilitate the reading of crowd emotion in a manner that is functionally related to the task at hand.

Our findings also provide new evidence that the processing that supports social decisions on crowd emotions are highly lateralized, in a manner that is relevant to the current task goal. Lateralized behavioral responses provide an opportunity to study the hemispheric asymmetries that enable cognitive functions. This hemispheric asymmetry enables flexible and adaptive processing optimized for the current task goal in dynamic environments (Rogers, Vallortigara, & Andrew, 2013), supporting the selection of appropriate, and inhibition of inappropriate, responses (Scott, 1962). This is particularly useful when a large number of complex stimuli (such as a crowd of emotional faces) and competing cognitive goals tax the processing capacity of the visual system, as was the case in our task.

The pattern of hemispheric lateralization in affective processing has been generally thought to rely on emotional valence (Craig, 2005; Davidson & Irwin, 1999), with aversive or negative stimuli lateralized to the right hemisphere (RH) and positive, approach-evoking stimuli lateralized to the left hemisphere (LH). However, we instead found that the lateralization effects in crowd emotion processing are actually goal-dependent, rather than being driven purely by stimulus valence. To our knowledge, this is the first demonstration that the same emotional stimuli can be biased differently in RH and LH, depending on the task goal and observers’ intent. Specifically, the current task goal (approach or avoidance) biased processing such that LVF/RH was superior for recognizing an angrier crowd during the avoidance task and for processing a happier crowd during the approach task. For RVF/LH, we found the opposite pattern: It was more accurate for implicit decisions, choosing a neutral crowd over a happier one during the avoidance task, and a neutral crowd over an angrier one during the approach task. Since the task-relevant emotions were anger and happiness for the avoidance and the approach tasks, respectively, our results suggest that LVF/RH is better at a clear, explicit decisions involving the task-congruent crowd emotion. Unlike the crowd emotion condition, however, we observed the overall RH advantage for the individual emotion condition in which emotional faces presented in the LVF led to better accuracy than in the RVF, consistent with previous findings (e.g., Borod et al., 1998; Dien, 2009; Hamilton & Vermeire, 1988; Kanwisher, McDermott, & Chun, 1997; Yovel, Tambini, & Brandman, 2008). Moreover, the accuracy was higher for angry faces than for happy faces in the LVF, indicating superior processing of angry faces over happy faces (Hansen & Hansen, 1988; Öhman, Lundqvist, & Esteves, 2001).

It is worth noting that the inference should be made very carefully from the divided visual field paradigm because it is a relatively indirect approach to localizing hemispheres with cognitive functions (see Ivry & Robertson, 1998). In particular, the interpretation of the results becomes very difficult when participants shift their gaze. To ensure that participants did not move their eyes while they performed the current task, we explicitly instructed participants to initiate each trial only after they fixated the central fixation cross. We also conducted a control eye tracking experiment where a new group of 18 participants performed the same task as in Experiment 1A, with their eye movement monitored throughout the experiment. We confirmed from this control experiment that the behavioral responses - both accuracy and RT - were comparable to our main results from Experiment 1A (SI.5) when their eye movement was monitored and restricted. Finally, we also replicated this hemispheric lateralization for crowd emotion processing in the three main behavioral experiments we report here and two replication experiments reported in the supplementary materials (SI.4). Thus, the laterality effect that we report here is robust to participants’ eye movements, different experimental settings, and with new cohorts of participants.

Different patterns of hemispheric lateralization for the crowd emotion vs. individual emotion processing suggest that they may rely on qualitatively distinct systems. Much behavioral evidence has been accumulated supporting this notion (e.g., Alvarez & Oliva, 2008; Ariely, 2001; Cant, Sun, & Xu, 2015; Chong et al., 2008; Chong & Evans, 2011; Chong & Treisman, 2005; Choo & Franconeri, 2010; Corbett & Oriet, 2011; Haberman, Brady, & Alvarez, 2015; Haberman & Whitney, 2010; Im & Halberda, 2013; Leib et al., 2012; Parkes et al., 2001, but see Myczek & Simons, 2008), although only a few recent fMRI studies have compared the neural representations of ensemble coding and individual processing (e.g., Cant & Xu, 2015, 2017; Huis in’t Veld & de Gelder, 2015). Cant and Xu (2015, 2017) showed that PPA and LO were preferentially engaged in texture perception and object processing, respectively; and Huis in’t Veld and de Gelder (2015) showed the greater anticipatory and action preparation activity in the areas including IPL, SPL, SFG, and premotor cortex for interactive body movement of a group of panicked people, compared to an unrelated movement of individuals. Because Cant and Xu (2015; 2017) used the stimuli of simple texture patches and objects and Huis in’t Veld and de Gelder (2015) removed the information about facial expression of people from their blurred video clips, the current study is the first to examine the distinct neutral substrates underlying the processing of facial crowds with varying emotional expressions and the processing individual faces and to show the evidence for distinct mechanisms supporting them.

The benefit of having distinct systems for ensemble coding and individual object processing is that these two processes can serve complementary functions. Global information extracted via ensemble coding influences processing of individual objects in many different ways. Because ensemble coding compresses properties of multiple objects into a compact description with a higher level of abstraction (Utochkin, 2015), it allows observers to surmount the severe limitations on individual object processing (Brady & Alvarez, 2015; Cohen, Dennett, & Kanwisher, 2016; Feigenson, 2011; Im & Chong, 2014; Im, Park, & Chong, 2015), imposed by attention or working memory (e.g., Cowan, 2001; Luck & Vogel, 1997; Pylyshyn & Storm, 1988; Treisman & Gelade, 1980). Furthermore, the global information of ensembles allows for an initial, rough analysis of visual inputs, which then biases and facilitates the processing of individual objects (Brady & Alvarez 2011; Haberman & Whitney, 2012; Im et al., 2015; Utochkin 2015). For example, extracted ensemble representation influences the individual object processing, by guiding detection of outliers in a set (e.g., pop-out visual search; Haberman & Whitney, 2012; Utochkin 2015), facilitating selection of an individual object at the center location of a set (e.g., Im et al., 2015), and biasing memory for individual objects towards the global mean (e.g., Brady & Alvarez, 2011). Our fMRI results showing differential activations for crowd emotion processing in the dorsal visual pathway vs. individual emotion processing in the ventral visual pathway have implications for such two distinct visual streams for global vs. local processing that can operate in parallel and interact with each other.

The results from the whole brain and the ROI analyses both support our hypothesis that the dorsal visual stream contributes to global processing for crowd emotion extraction whereas the ventral visual stream contributes more to object-based processing of emotion in individual faces. The ROIs that we reported here are the areas that have been found in prior work as supporting the input predominantly from the M-dominant dorsal pathway (IPS and SFG; Courtney et al., 1996; Courtney et al., 1998; Sala & Coutney, 2007; Takahashi et al., 2013; Wilson et al., 1993) and from the P-dominant ventral pathway (FG; Denys et al., 2004; Grill-Spector & Malach, 2004; Haxby et al., 1991; Kanwisher et al., 1997; Purves et al., 2004; Takahashi et al., 2013; Taylor et al., 2007). Moreover, we also observed that bilateral IPS and SFG were activated by M-pathway stimuli and bilateral FG was activated by P-pathway stimuli in our pilot study using functional localizer scans for M-pathway and P-pathway regions (SI.6). In this pilot study, we used the sinusoidal counter-phase flickers biasing M-pathway (a low-luminance contrast, black-white grating with a low spatial frequency and 15 Hz flicker) and biasing P-pathway (an isoluminant, high color-contrast red-green grating with high spatial frequency with slow, 5 Hz flicker), following the method detailed in Denison et al. (2014). Although different types of stimuli (face stimuli in the current study vs. gratings in the pilot study) and different cohorts of participants were employed, the localizer scans showed the activations in the adjacent foci to our ROIs. This confirms that our ROIs selection is reasonable, effectively reflecting the differential contribution of M- and P-pathways to neural processing of the crowd emotion and individual emotion. Our previous (Kveraga et al., 2007; Kveraga et al., 2007; Kveraga, 2014; Thomas et al., 2012) and current (Adams et al., in prep.; Im et al., in prep.) studies using object, letter, scene, and face stimuli biased towards M or P pathways likewise support this dorsal/ventral split in activation for M and P stimuli, respectively.

Processing of crowd emotion appears to be achieved in a global, parallel fashion, rather than serially for the following reasons. First of all, we found that the RTs for the crowd emotion condition were equivalent to those for the individual emotion, despite many more faces that needed to be processed in the crowd condition compared to the individual face condition. Second, simply sampling any one or two individual faces from each crowd in our stimuli would lead to the chance level of accuracy, because a half of the individual members in the neutral crowd were always more intense than a half of the angry crowd in our stimuli. Thus, sampling faces with an extreme emotional expression can not be an explanation for our findings. Third, the participants were equally accurate and fast when they viewed the facial crowds containing 8 faces or 12 faces. Therefore, we suggest that extracting crowd emotion relies on a parallel, global process, rather than on a sequential sampling of individual members (e.g., Myczek & Simons, 2008). In various feature dimensions, the notion of global averaging has been previously tested, by using empirical approaches showing that multiple stimuli were integrated (e.g., Leib et al., 2014), ideal observer analysis (e.g., Haberman et al., 2009; Im & Halberda, 2013), equivalent noise (e.g., Florey et al., 2016), or general linear modeling (e.g., Hubert-Wallander & Boynton, 2015). Consistent with this prior work, the current findings suggest that people do average different facial expressions to make social decisions about facial crowds, and such ensemble coding of crowds of faces is achieved via a distinct mechanism from that supporting individual object processing.

To conclude, here we have reported evidence for distinct mechanisms dedicated to processing of crowd emotion and individual face emotion, which are biased towards different visual streams (dorsal vs. ventral), and show different patterns of hemispheric lateralization. The differential engagement of the dorsal stream regions and the complementary functions of the left and right hemispheres both suggest that processing of crowd emotion is specialized for action execution that is highly flexible and goal-driven, allowing us to trigger a rapid and appropriate reaction to our social environment. Furthermore, we have shown that observers’ goals - to avoid or approach - can exert powerful influences on the perception accuracy of crowd emotion, highlighting the importance of understanding the interplay of ensemble coding of crowd emotion and social vision.

## General Methods

### Participants

In Experiment 1, a total of 42 undergraduate students participated: 21 subjects (12 female) participated in the avoidance task (Exp.1A) and a different cohort of 21 participants (11 female) participated in the approach task (Exp.1B). No subjects were excluded from the behavioral data analysis. In Experiment 2, a new group of 32 (18 female) undergraduate students participated. Two participants were excluded from further analyses because they made too many late responses (e.g., RTs longer than 2.5s). Thus, the behavioral and fMRI analyses for the Experiment 2 were done with a sample of 30 participants. All the participants had normal color vision and normal or corrected-to-normal visual acuity. Their informed written consent was obtained according to the procedures of the Institutional Review Board at the Pennsylvania State University. The participants received monetary compensation or a course credit.

### Apparatus and stimuli

Stimuli were generated with MATLAB and Psychophysics Toolbox (Brainard, 1997; Pelli, 1997). In each crowd stimulus (Fig. 1A), either 4 or 6 morphed faces were randomly positioned in each visual field (right and left) on a grey background. Therefore, our facial crowd stimuli comprised either 8 or 12 faces. We used a face-morphing software (Norrkross MorphX) to create a set of 51 morphed faces from two highly intense, prototypical facial expressions of the same person for a set of six different identities (3 male and 3 female faces), taken from the Ekman face set (Ekman & Friesen, 1976). The morphed face images were controlled for luminance, and the emotional expression of the faces ranged from happy to angry (Fig. 1B), with 0 in Emotional Unit (EU) being neutral (morph of 50% happy and 50% angry), +25 in EU being the happiest (100% happy), and -25 in EU being the angriest (100% angry). Because the morphed face images were linearly interpolated (in 2% increments) between two extreme faces, they were separated from one another by EU of intensity such that Face 1 was one EU happier than Face 2, and so on. Therefore, the larger the separation between any two morphed faces in EU, the easier it was to discriminate them. Such morphing approach was adapted from the previous studies on ensemble coding of faces (e.g., Haberman & Whitney, 2007).

Since the previous literature on averaging of other visual features showed that the range of variation is an important determinant of averaging performance (e.g., size or hue: Maule & Franklin, 2015; Utochkin & Tiurina, 2014), we kept the range of faces the same (i.e., 18 in emotional units) across the two set sizes. One of the two crowds in either left or right visual field always had the mean value of zero in emotional units, which is neutral on average, and the other had the emotional mean of +9 (very happy; morphing of angry 32 % and happy 68%), +5 (somewhat happy; morphing of angry 40% and happy 60%), -9 (very angry; morphing of angry 68% and happy 32%), and -5 (somewhat angry; morphing of angry 60% and happy 40%). Thus, the sign of such offset between the emotional and neutral crowds in EU indicates the valence of the emotional crowd to compared to the neutral: The positive values indicate more positive (happier) crowd emotion compared with the neutral and the negative values indicate more negative (angrier) mean emotion.

In order to avoid the possibility that participants simply sampled one or two single faces from each set and compare them to do the crowd emotion task, we ensured that 50% of the individual faces in the neutral set were more expressive than 50% of the individual faces in the emotional sets to be compared. For example, half of the members of the neutral set were angrier than a half of the members of the angry crowd. This manipulation allowed us to assess whether participants used such ‘sampling strategy (Myczek & Simons, 2008)’ rather than extracting average, because sampling one or two members in a set would yield 50% of accuracy in this setting.

Stimuli for the individual emotion condition (Fig. 1C; only included in the fMRI study) comprised one emotional face (either angry or happy) and one neutral face from the same set of morphed face images randomly positioned in the same invisible frame surrounding the crowd stimuli in each visual field. The offsets between the emotional and neutral faces remained the same as those in facial crowd stimuli. To ensure that the difference is not due to the confound of simply having more ‘stuff’ in crowd emotion condition, compared to the individual emotion condition, we included scrambled faces in the individual emotion condition so that the same number of the face-like blobs were presented as in the crowd emotion condition. This ensured that any differences are not due to low-level visual differences in the stimulus displays, but rather to how many resolvable emotional faces participants had to discriminate on each trial (2 vs. 8 or 12).

On one half of the trials, the emotional stimulus (i.e., happy or angry: ± 5 or ± 9 EU away from the mean) was presented in the left visual field and the neutral stimulus was presented in the right visual field, and it was switched for the other half of the trials. Each face image subtended 2° x 2° of visual angle, and face images were randomly positioned within an invisible frame subtending 13.29° x 18.29°, each in the left and right visual fields. The distance between the proximal edges of the invisible frames in left and right visual fields was 3.70°.

### Procedure

Participants in Experiment 1 sat in a chair at individual cubicles about 61 cm away from a computer with a 48 cm diagonal screen (refresh rate = 60 Hz). Participants in Experiment 2 were presented with the stimuli rear-projected onto a mirror attached to a 64-channel head coil in the fMRI scanner. Fig. 1A illustrates a sample trial of the experiment. Participants were presented with visual stimuli for 1 second, followed by a blank screen for 1.5 second. The participants were instructed to make a key press as soon as possible to indicate which of the two crowds of faces or two single faces on the left or right they would rather avoid. They were explicitly informed that the correct answer was to choose either the crowd or the face showing a more negative (e.g., angrier) emotion for the avoidance task and a more positive (e.g., happier) emotion for the approach task. Responses that were made after 2.5 seconds were considered late and excluded from data analyses. Feedback for correct, incorrect, or late responses was provided after each response.

In Experiment 1, half of the participants performed the avoidance task and the other half performed the approach task. Experiment 1 had a 4 (emotional distance between facial crowds, -9, -5, 5, or 9) x 2 (visual field of presentation, LVF and RVF) x 2 (set size: 4 or 6 faces in each visual field) design, and the sequence of total 320 trials (20 repetitions per condition) was randomized. In Experiment 2 (fMRI), all the participants performed the avoidance task. Because we needed more trials for statistical power for fMRI data analyses and we observed no effect by the number of crowd members on crowd emotion perception (Fig.S1), we only used crowd stimuli containing 4 faces in Experiment 2. Thus, Experiment 2 had a 2 (stimulus type: crowd and individual) x 4 (emotional distance) x 2 (visual field of presentation) design and the sequence of total 512 trials (32 repetitions per conditions) was optimized for hemodynamic response estimation efficiency using the *optseq2* software (https://surfer.nmr.mgh.harvard.edu/optseq/).

### fMRI data acquisition and analysis

fMRI images of brain activity were acquired using a 3 T scanner (Siemens Magnetom Prisma) located at The Pennsylvania State University Social, Life, and Engineering Sciences Imaging Center. High resolution anatomical MRI data were acquired using T1-weighted images for the reconstruction of each subject’s cortical surface (TR = 2300 ms, TE = 2.28 ms, flip angle = 8°, FoV = 256 x 256 mm^2^, slice thickness = 1 mm, sagittal orientation). The functional scans were acquired using gradient-echo EPI with a TR of 2000 ms, TE of 28ms, flip angle of 52° and 64 interleaved slices (3 x 3 x 2 mm). Scanning parameters were optimized by manual shimming of the gradients to fit the brain anatomy of each subject, and tilting the slice prescription anteriorly 20-30° up from the AC-PC line as described in the previous studies (Deichmann et al., 2003; Kveraga et al., 2007; Wall, Walker, & Smith, 2009), to improve signal and minimize susceptibility artifacts in the brain regions including OFC and amygdala (Kringelbach & Rolls, 2004). We acquired 780 functional volumes per subject in four functional runs, each lasting 6.5 min.

The acquired fMRI mages were pre-processed using SPM8 (Wellcome Department of Cognitive Neurology). The functional images were corrected for differences in slice timing, realigned, corrected for movement-related artifacts, coregistered with each participant’s anatomical data, normalized to the Montreal Neurological Institute template, and spatially smoothed using an isotropic 8-mm full width half-maximum Gaussian kernel. Outliers due to movement or signal from preprocessed files, using thresholds of 3 SD from the mean, 0.75 mm for translation and 0.02 radians rotation, were removed from the data sets, using the ArtRepair software (Mazaika et al., 2009). Subject-specific contrasts were estimated using a fixed-effects model. These contrast images were used to obtain subject-specific estimates for each effect. For group analysis, these estimates were then entered into a second-level analysis treating participants as a random effect, using one-sample t-tests at each voxel. The resulting contrasts were thresholded at *p* < 0.001 (uncorrected) and a minimal cluster size of 10 voxels. For visualization and anatomical labeling purposes, all group contrast images were overlaid onto the inflated group average brain, by using 2D surface alignment techniques implemented in FreeSurfer (Fischl et al. 2004).

For the region of interest (ROI) analyses, we extracted the BOLD activity from IPS, SFG, FG. We defined a contrast between all the visual stimulation trials vs. background (Null trials). From this contrast, we localized each of the ROIs based on the peak activation within the anatomical label obtained by the anatomical parcellation of the normalized brain (Tzourio-Mazoyer et al., 2002). The [x y z] coordinates for these ROIs were [30 -70 42] for IPS, [36 6 58] for SFG, and [33 -58 -10] for FG. The coordinates for IPS and SFG were adjacent to the regions that have been reported in the previous study as showing the robust intra-stream connectivity (dorsal visual stream; Takahashi et al., 2013). Moreover, the coordinate for the FG was has been also localized as the right FFA (Fusiform Face Area) in the activation maps by Spiridon, Fischl, & Kanwisher (2006). The beta weights were extracted for crowd emotion and individual emotion conditions using the rfxplot toolbox (http://rfxplot.sourceforge.net) for SPM. Around the [x y z] coordinate for each of our ROIs, we defined a 6mm sphere around it. Using the rfxplot toolbox in SPM8, we extracted all the voxels from each individual participant’s functional data within that sphere. The extracted beta weights for each of the four trial conditions were subjected to repeated-measures ANOVA and to correlation analyses with behavioral accuracy measurements.

## Footnotes

1 We also conducted RT analyses using each participant’s median RT. Just as mean RT, median RT was significantly slower for the avoidance task than for the approach task (1.16 second vs. 0.97 second on average: *t*(40) = 1.995, *p* < 0.05, Cohen’s d = 0.632). We also confirmed that median RTs yielded the same results for all the other findings reported in this manuscript (see Supplementary Information).

## Acknowledgments

This work was supported by the National Institutes of Health R01MH101194 to K.K. and to R.B.A., Jr.

Kestas Kveraga: Reginald B. Adams, Jr.

Data collection was conducted at the Pennsylvania State University. Informed written consent was obtained in all studies according to the procedures of the Institutional Review Board at the Pennsylvania State University. The participants received a course credit for their participation.

## Author Contributions

H. Y. Im, R. B. Adams, and K. Kveraga developed the study concept, and all authors contributed to the study design. Testing and data collection were performed by H.Y. Im, C. A. Cushing, T. G. Steiner, and D. N. Albohn. H. Y. Im analyzed the data and all the authors wrote the manuscript.

## References

Adams, R.B., Jr., Ambady, N., Macrae, C. N., & Kleck, R. E. (2006). Emotional expressions forecast approach-avoidance behavior. Motivation & Emotion, 30, 179–188.

Adams, R. B., Jr., Hess, U., & Kleck, R. E. (2015). The Intersection of Gender-Related Facial Appearance and Facial Displays of Emotion. Emotion Review, 7, 5–13.

Adams, R. B., Jr., & Kveraga, K. (2015). Social vision: Functional forecasting and the integration of compound social cues. Review of Philosophy and Psychology, 6, 591–610.

Alvarez, G. A. (2011). Representing multiple objects as an ensemble enhances visual cognition. Trends in Cognitive Sciences, 15, 3, 122–131.

Alvarez, G. A., & Oliva, A. (2008). The representation of simple ensemble visual features outside the focus of attention. Psychological Science, 19, 392–398.

Ariely, D. (2001). Seeing sets: Representation by statistical properties. Psychological Science, 12, 157–162.

Bargh, J. A., Chen, M., & Burrows, L. (1996). Automaticity of social behavior: Direct effects of trait construct and stereotype activation on action. Journal of Personality & Social Psychology, 71, 230–244.

Becker, D. V., Kenrick, D. T., Neuberg, S. L., Blackwell, K. C., & Smith, D. M. (2007). The confounded nature of angry men and happy women. Journal of Personality and Social Psychology, 92, 179–190.

Borod, J. C., Cicero, B. A., Obler, L. K., Welkowitz, J., Erhan, H. M., Santschi, C., Grunwald, I. S., Agosti, R. M., & Whalen, J. R. (1998). Right hemisphere emotional perception: evidence across multiple channels. Neuropsychology, 12, 446–458.

Brady, T. F., & Alvarez, G. A. (2011). Hierarchical encoding in visual working memory: ensemble statistics bias memory for individual items. Psychological Science, 22, 384–392.

Brady, T. F., & Alvarez, G. A. (2015). No evidence for a fixed object limit in working memory: Ensemble representations inflate estimates of working memory capacity for complex objects. Journal of Experimental Psychology: Learning, Memory and Cognition, 41, 921–929.

Brainard, D. H. (1997). The Psychophysics Toolbox. Spatial vision, 10, 433–436.

Brunyé, T. T., Howe, J. L., & Mahoney, C. R. (2014). Seeing the crowd for the bomber: Spontaneous threat perception from static and randomly moving crowd simulations. Journal of Experimental Psychology: Applied, 20, 303–322.

Cant, J. S., Sun, S. Z., & Xu, Y. (2015). Distinct cognitive mechanisms involved in the processing of single objects and object ensembles. Journal of Vision, 15, 1–21.

Cant, J. S. & Xu, Y. (2015). The impact of density and ratio on object-ensemble representation in human anterior-medial ventral visual cortex. Cerebral Cortex, 25, 4226–4239.

Cant, J. S. & Xu, Y. (2017). The contribution of object shape and surface properties to object-ensemble representation in anterior-medial ventral visual cortex. Journal of Cognitive Neuroscience, 29, 398–412.

Chong, S. C., & Treisman, A. (2003). Representation of statistical properties. Vision Research, 43, 393–404.

Choo, H., & Franconeri, S. L. (2010). Objects with reduced visibility still contribute to size averaging. Attention, Perception & Psychophysics, 72, 86–99.

Chong, S. C., & Evans, K. K. (2011). Distributed vs. Focused attention (count vs. estimate). Wiley Interdisciplinary Reviews: Cognitive Science, 2, 634–638.

Chong, S. C., & Treisman, A. (2005). Attentional spread in the statistical processing of visual displays. Perception & Psychophysics, 67, 1–13.

Chong, S. C., Joo, S. J., Emmmanouil, T., & Treisman, A. (2008). Statistical processing: Not so implausible after all. Perception & Psychophysics, 70, 1327–1334.

Cohen, M.A., Dennett, D. C., & Kanwisher, N. (2016). What is the bandwidth of perceptual experience? Trends in Cognitive Sciences, 19, 324–335.

Corbett, J. E., & Oriet, C. (2011). The whole is indeed more than the sum of its parts: Perceptual averaging in the absence of individual item representation. Acta Psychologica, 138, 289–301.

Courtney, S. M., Ungerleider, L. G., Keil, K., & Haxby, J. V. (1996). Object and spatial visual working memory activate separate neural systems in human cortex. Cerebral Cortex, 6, 39–49.

Courtney, S. M., Petit, L., Maisog, J. M., Ungerleider, L. G., & Haxby, J. V. (1998). An area specialized for spatial working memory in human frontal cortex. Science, 279, 1347–1351.

Cowan, N. (2001). The magical number 4 in short-term memory: A reconsideration of mental storage capacity. Behavioral and Brain Sciences, 24, 87–185.

Craig, A. D. (2005). Forebrain emotional asymmetry: a neuroanatomical basis? Trends in Cognitive Sciences, 9, 566–571.

Dakin, S. C., & Watt, R. J. (1997). The computation of orientation statistics from visual texture. Vision Research, 37, 3181–3192.

Davidson, R. J. (1992). Anterior cerebral asymmetry and the nature of emotion. Brain & Cognition, 20, 125–151.

Davidson, R. J. (1995). Brain Asymmetry, Cerebral asymmetry, emotion, and affective style. In: Davidson, R.J., Hugdahl, K, editors (MIT Press, Cambridge, MA), pp. 361–387.

Davidson, R. J., & Irwin, W. (1999). The functional neuroanatomy of emotion and affective style. Trends in Cognitive Sciences, 3, 11–21.

De Fockert, J. W., & Wolfenstein, C. (2009). Rapid extraction of mean identity from sets of faces. Quarterly Journal of Experimental Psychology, 62, 1716–1722.

De Renzi, E. (1986). Prosopagnosia in two patients with CT scan evidence of damage confined to the right hemisphere. Neuropsychologia, 24, 385–389.

Deichmann, R., Gottfried, J. A., Hutton, C., & Turner, R. (2003). Optimized EPI for fMRI studies of the orbitofrontal cortex. Neuroimage, 19, 430–441.

Denison, R. N., Vu, A. T., Yacoub, E., Feinberg, D. A., & Silver, M. A. (2014). Functional mapping of the magnocellular and parvocellular subdivisions of human LGN. Neuroimage, 102, 358–369.

Denys, K., Vanduffel, W., Fize, D., Nelissen, K., Peuskens, H., Van Essen, D., & Orban, G. A. (2004). The processing of visual shape in the cerebral cortex of human and nonhuman primates: a functional magnetic resonance imaging study. Journal of Neuroscience, 24, 2551–2565.

Dien, J. (2009). A tale of two recognition systems: implications of the fusiform face area and the visual word form area for lateralized object recognition models. Neuropsychologia, 47, 116.

Ekman, P., & Friesen, W. V. (1976). Pictures of facial affect. Consulting Psychologists Press; Palo Alto, CA.

Elliot, A. J. (1999). Approach and avoidance motivation and achievement goals. Educational Psychologist, 34, 169–189.

Fabes, R. A., & Martin, C. L. (1991). Gender and age stereotypes of emotionality. Personality and Social Psychology Bulletin, 17, 532–540.

Feigenson, L. (2011). Objects, sets, and ensemble. In S. Dehaene & E. Brannon (Eds.), Space, Time, and Number in the Brain: searching for the Foundations of Mathematical Thought. London: Elsevier.

Fischer, J., & Whitney, D. (2011). Object-level visual information gets through the bottleneck of crowding. Journal of Neurophysiology, 106, 1389–1398.

Fischl, B., Van Der Kouwe, A., Destrieux, C., et al. (2004). Automatically parcellating the human cerebral cortex. Cerebral Cortex, 14, 11–22.

Fischl, B., van der Kouwe, A., Destrieux, C., Halgren, E., Ségonne, F., Salat, D. H., Busa, E., Seidman, L. J., Goldstein, J., Kennedy, D., Caviness, V., Makris, N., Rosen, B., & Dale, A. M. (2004). Automatically parcellating the human cerebral cortex. Cerebral Cortex, 14, 11–22.

Florey, J., Clifford, C. W., Dakin, S., & Mareschal, I. (2016). Spatial limitations in averaging social cues. Scientific Reports, 6, 32210.

Freud, E., Plaut, D. C., & Behrmann, M. (2016). ‘What’ is happening in the dorsal visual pathway. Trends in Cognitive Sciences, 20, 773–784.

Goodale, M. A., & Milner, A. D. (1992). Separate visual pathways for perception and action. Trends in Neurosciences, 15, 20–25.

Grill-Spector, K., & Malach, R. (2004). The human visual cortex. Annual Review of Neuroscience, 27, 649–677.

Haberman, J., & Whitney, D. (2010). The visual system discounts emotional deviants when extracting average expression. Attention, Perception, & Psychophysics, 72, 1825–1838.

Haberman, J., & Whitney, D. (2007). Rapid extraction of mean emotion and gender from sets of faces. Current Biology, 17, R751–R753.

Haberman, J. & Whitney, D. (2011). Efficient summary statistical representation when change localization fails. Psychonomic Bulletin and Review, 18, 855–859.

Haberman, J., & Whitney, D. (2012). ‘Ensemble Perception: Summarizing the scene and broadening the limits of visual processing.’ In J. Wolfe and L. Robertson (Eds.), From Perception to Consciousness: Searching with Anne Treisman. Oxford University Press.

Haberman, J., Brady, T.F., & Alvarez, G.A. (2015). Individual differences in ensemble perception reveal multiple, independent levels of ensemble representation. Journal of Experimental Psychology: General, 144, 432–446.

Haberman, J., Harp, T., & Whitney, D. (2009). Averaging facial expression over time. Journal of Vision, 9, 1–13.

Halberda, J., Sires, S. F., & Feigenson, L. (2006). Multiple spatially overlapping sets can be enumerated in parallel. Psychological Science, 17, 572–576.

Hamilton, C. R., & Vermeire, B. A. (1988). Complementary hemispheric specialization in monkeys. Science, 242, 1691–1694.

Hansen, C. H., & Hansen, R. D. (1988). Finding the face in the crowd: An anger superiority effect. Journal of Personality and Social Psychology, 54, 917–924.

Haxby, J. V., Grady, C. L., Horwitz, B., et al. (1991). Dissociation of object and spatial visual processing pathways in human extrastriate cortex. Proceedings of the National Academy of Science, USA, 88, 1621–1625.

Horstmann, G. (2003). What do facial expressions convey: Feeling states, behavioral intentions, or action requests? Emotion, 3, 150–166.

Hubert-Wallander, B., & Boynton, G. M. (2015). Not all summary statistics are made equal: Evidence from extracting summaries across time. Journal of Vision, 15, 1–12.

Huis in ‘t Veld, E. M. J., & de Gelder, B. (2015). From individual fear to mass panic. The neurological basis of crowd perception. Human Brain Mapping, 36, 2338–1351.

Im, H. Y., & Chong, S. C. (2014). Mean size as a unit of visual working memory. Perception, 43, 663–676.

Im, H. Y., & Halberda, J. (2013). The effects of sampling and internal noise on the representation of ensemble average size. Attention, Perception, & Psychophysics, 75, 278–286.

Im, H. Y., Park, W. J., & Chong, S. C. (2015). Ensemble statistics as units of selection. Journal of Cognitive Psychology, 27, 114–127.

Ivry, R. B., & Robertson, L. C. (1998). The two sides of perception. Cambridge (MA): MIT Press.

Ji, L., Chen, W., & Fu, X. (2014). Different roles of foveal and extrafoveal vision in ensemble representation for facial expressions. Engineering Psychology and Cognitive Ergonomics Lecture Notes in Computer Science, 8532, 164–173.

Kanwisher, N., McDermott, J., & Chun, M. M. (1997). The fusiform face area: A module in human extrastriate cortex specialized for face perception. Journal of Neuroscience, 17, 4302–4311.

Kringelbach, M. L., & Rolls, E. T. (2004). The functional neuroanatomy of the human orbitofrontal cortex: evidence from neuroimaging and neuropsychology. Progress in Neurobiology, 72, 341–372.

Kveraga, K., Boshyan, J., & Bar, M. (2007). The magnocellular trigger of top-down facilitation in object recognition. Journal of Neuroscience, 27, 13232–13240.

Kveraga, K., Ghuman, A. S., & Bar, M. (2007). Top-down predictions in the cognitive brain. Brain and Cognition, 65, 145–168.

Kveraga, K. (2014). Threat perception in visual scenes: dimensions, action and neural dynamics. In K. Kveraga & M. Bar (Eds.) Scene Vision: making sense of what we see. Cambridge, MIT Press, pp. 291–307.

Leib, A. Y., Puri, A. M., Fischer, J., Bentin, S., Whitney, D., & Robertson, L. (2012). Crowd perception in prosopagnosia. Neuropsychologia, 50, 1698–1707.

Leib, A. Y., Fischer, J., Liu, Y., Qiu, S., Robertson, L., & Whitney, D. (2014). Ensemble crowd perception: A viewpoint-invariant mechanism to represent average crowd identity. Journal of Vision, 14, 1–13.

Livingstone, M. S., & Hubel, D. E. (1988). Segregation of form, color, movement, and depth: anatomy, physiology, and perception. Science, 240, 740–749.

Luck, S., & Vogel, E. K. (1997). The capacity of visual working memory for features and conjunctions. Nature, 390, 279–281.

Marsh, A. A., Ambady, N., & Kleck, R. E. (2005). The effects of fear and anger facial expressions on approach- and avoidance-related behaviors. Emotion, 5, 119–124.

Maule, J., & Franklin, A. (2015). Effects of ensemble complexity and perceptual similarity on rapid averaging of hue. Journal of Vision, 15, 1–18.

Mazaika, P. K., Hoeft, F., Glover, G. H., & Reiss, A. L. (2009). Methods and software for fMRI analysis for clinical subjects, in Poster Session Presented at the Meeting of Human Brain Mapping 2009 (San Fransisco, CA)

Merigan, W. H., & Maunsell, J. H. (1993). How parallel are the primate visual pathways? Annual Review of Neuroscience, 16, 369–402.

Milner, A. D., & Goodale, M.A. (1995). The Visual Brain in Action. Oxford University Press, Oxford.

Milner, A. D., & Goodale, M. A. (2008). Two visual systems re-viewed. Neuropsychologia, 46, 774–785.

Myczek, K., & Simons, D. J. (2008). Better than average: Alternatives to statistical summary representations for rapid judgments of average size. Perception & Psychophysics, 70, 772–788.

Neumann, M. F., Schweinberger, S. R., & Burton, A. M. (2013). Viewers extract mean and individual identity from sets of famous faces. Cognition, 128, 56–63.

Öhman, A., Lundqvist, D., & Esteves, F. (2001). The face in the crowd revisited: A threat advantage with schematic stimuli. Journal of Personality and Social Psychology, 80, 381–396.

Parkes, L., Lund, J., Angelucci, A., Solomon, J. A., & Morgan, M. (2001). Compulsory averaging of crowded orientation signals in human vision. Nature Neuroscience, 4, 739–744.

Pelli, D. G. (1997). The VideoToolbox software for visual psychophysics: transforming numbers into movies. Spatial vision, 10, 437–442.

Purves, D., Augustine, G. J., Fitzpatrick, D., Hall, W. C., LaMantia, A-S., McNamara, J. O., & Williams, S. M. (2004). Neuroscience (3rd ed.). Massachusetts U.S.A.: Sinauer Associates Inc.

Pylyshyn, Z. W., & Storm, R. W. (1988). Tracking multiple independent targets: Evidence for a parallel tracking mechanism. Spatial Vision, 3, 179–197.

Rogers, L. J., Vallortigara, G., & Andrew, R. J. (2013). Divided brains: The biology and behavior of brain asymmetries. Cambridge, UK: Cambridge University Press.

Sala, J. B., & Courtney, S. M. (2007). Binding of what and where during working memory maintenance. Cortex, 43, 5–21.

Sawatari, A., & Callaway, E. M. (1996). Convergence of magno- and parvocellular pathways in layer 4B of macaque primary visual cortex. Nature, 380, 442–446.

Schiller, P. H., & Logothetis, N. K. (1990). The color- opponent and broad-band channels of the primate visual system. Trends in Neuroscience, 13, 392–398.

Scott, W. A. (1962). Cognitive complexity and cognitive flexibility. American Sociological Association, 25, 405–414.

Silberman, E. K., & Weingartner, H. (1986). ‘Hemispheric lateralization of functions related to emotion’. Brain and Cognition, 5, 322–353.

Spiridon, M., Fischl, B., & Kanwisher, N. (2006). Location and spatial profile of category-specific regions in human extrastriate cortex. Human Brain Mapping, 27, 77–89.

Sweeny, T. D., & Whitney, D. (2014). Perceiving crowd attention: Ensemble perception of a crowd’s gaze. Psychological Science, 25, 1903–1913.

Sweeny, T. D., Haroz, S., & Whitney, D. (2013). Perceiving group behavior: Sensitive ensemble coding mechanisms for biological motion of human crowds. Journal of Experimental Psychology: Human Perception and Performance, 39, 329–337.

Takahashi, E., Ohki, K., & Kim, D-S. (2013). Dissociation and convergence of the dorsal and ventral visual streams in the human prefrontal cortex. Neuroimage, 65, 488–498.

Taylor, J. C., Wiggett, A. J., & Downing, P. E. (2007). Functional MRI analysis of body and body part representations in extrastriate and fusiform body parts. Journal of Neurophysiology, 98, 1626–1633.

Thomas, C., Kveraga, K., Huberle, E., Karnath, H-O., & Bar, M. (2012). Enabling global processing in simultanagnosia by psychophysical biasing of visual pathways. Brain, 135, 1578–1585.

Treisman, A. M., & Gelade, G. (1980). A feature-integration theory of attention. Cognitive Psychology, 12, 97–136.

Tzourio-Mazoyer, N., Landeau, B., Papathanassiou, D., Crivello, F., Etard, O., Delcroix, N., Mazoyer, B., & Joliot, M. (2002). Automatic anatomical labelling of activations in SPM using a macroscopic anatomical parcellation of the MNI MRI single-subject brain. Neuroimage, 15, 273–289.

Utochkin, I. S., & Tiurina, N. A. (2014). Parallel averaging of size is possible but range-limited: a reply to Marchant, Simons, and De Fockert. Acta Psychologica, 146, 7–18.

Utochkin, I. S. (2015). Ensemble summary statistics as a basis for rapid visual categorization. Journal of Vision, 15, 1–14.

Vuilleumier, P., Armony, J. L., Driver, J., & Dolan, R. J. (2003). Distinct spatial frequency sensitivities for processing faces and emotional expressions. Nature Neuroscience, 6, 624–631.

Wada, Y., & Yamamoto, T. (2001). Selective impairment of facial recognition due to a haematoma restricted to the right fusiform and lateral occipital region. Journal of Neurology, Neurosurgery, and Psychiatry, 71, 254–257.

Wall, M. B., Walker, R., & Smith, A. T. (2009). Functional imaging of the human superior colliculus: an optimised approach. Neuroimage, 47, 1620–1627.

Watamaniuk, S. N. J., & Sekuler, R. (1992). Temporal and spatial integration in dynamic random-dot stimuli. Vision Research, 32, 2341–2347.

Wilson, F. A., Scalaidhe, S. P., & Goldman-Rakic, P. S. (1993). Dissociation of object and spatial processing domains in primate prefrontal cortex. Science, 260, 1955–1958.

Winston, J. S., Vuilleumier, P., & Dolan, R. J. (2003). Effects of low-spatial frequency components of fearful faces on fusiform cortex activity. Current Biology, 13, 1824–1829.

Yang, J.-W., Yoon, K. L., Chong, S. C., & Oh, K. J. (2013). Accurate but pathological: Social anxiety and ensemble coding of emotion. Cognitive Therapy and Research, 37, 572–578.

Yovel, G., Tambini, A., & Brandman, T. (2008). The asymmetry of the fusiform face area is a stable individual characteristic that underlies the left-visual-field superiority for faces. Neuropsychologia, 46, 3061–3068.

